# Encoding visual stimuli by striatal neurons

**DOI:** 10.1101/2025.09.15.676378

**Authors:** Job Perez-Becerra, Ricardo Velazquez-Contreras, Luis A. Tellez, Luis Carrillo-Reid

## Abstract

To move through the world, animals extract visual features from objects. Although visual object encoding is considered a cortical attribute, subcortical areas also contain visual processing circuits. Using electrophysiological recordings of spiny projection neurons (SPNs) located in a target region of the primary visual cortex, we identified striatal modules that process basic visual features. SPNs from the direct and indirect pathways exhibited high orientation selectivity in the absence of behavioral contingencies. SPNs population activity reliably predicted different visual features. Pathway-specific silencing of SPNs revealed that high orientation selectivity depends on intra-striatal connectivity. Thus, specialized striatal modules are active participants in visual feature extraction. These findings indicate parallel visual information processing in basal ganglia circuits, expanding current models of visual perception beyond cortical networks.

## Introduction

For more than 60 years, the study of visual information processing has focused on hierarchically organized visual cortical areas with the goal of understanding the role of different cortices in visual perception. It is known that a major function of primary visual cortex (V1) is to elaborate orientation selectivity (*1, 2*). In mice, orientation selectivity is encoded by V1 cortical neurons (*3–5*). However, it is unknown if subcortical neurons also show orientation selectivity. As the gateway of basal ganglia (BG), the striatum is considered a hub that oversees the integration of contextual and motor information from several cortical areas to optimize the execution of behaviorally relevant actions (*6–9*). Thus, striatal neurons are thought to process multiplexed signals to guide movements (*10, 11*).

Early studies in monkeys, described that striatal responses to visual stimuli were unreliable and mainly related to behavioral contingencies rather than physical characteristics of sensory stimuli (*12–16*). Consequently, for decades striatal visual responses have been deemed as the result of processed visual information related to learned behaviors (*17–20*) dismissing the role of striatal neuronal activity in processing basic visual features.

Using computational and neuroanatomical approaches, it has been shown that a specific region of the dorsomedial striatum (DMS) receives direct projections from primary visual cortex (*21*). Moreover, optogenetic activation of V1 axons projecting to DMS demonstrated functional excitatory synapses onto striatal neurons from the direct and indirect BG pathways (*22*) suggesting that topographically localized striatal subpopulations could encode similar features of visual stimuli as V1. However, how BG structures process basic features of visual stimuli remains poorly understood (*23, 24*).

## Results

### Responses to visual stimuli in neurons from the dorsomedial striatum

Although striatal visual responses have been studied during learned behaviors or multisensory integration in rodents (*25–27*), how dorsomedial striatal neurons process specific visual features conveyed by V1 projections remains unknown.

To characterize striatal responses to visual stimuli in the absence of behavioral conditioning we performed electrophysiological recordings with silicon probes in untrained awake head-fixed mice (Figure 1A) in the dorsomedial striatal region that is adjacent to the dorsal half of the lateral ventricle and receives direct projections from primary visual cortex (*21*). To isolate sensory responses from motor-related activity, we analyzed neuronal responses to visual stimuli during motionless periods (741 neurons from 24 mice). In these conditions we observed increased responses in ~55% of the recorded neurons even in the absence of behavioral conditioning (Figure 1B). 74% of the recorded neurons were putative spiny projection neurons (SPNs) that can be reliably distinguished from putative fast spiking interneurons (FSIs) (*28*). Significantly increased firing rates were observed in 319 SPNs and 91 FSIs. From the remaining neurons, 195 SPNs and 80 FSIs were unresponsive to visual stimulation, whereas 33 SPNs and 23 FSIs decreased their firing rates after the onset of visual stimulation (Figure 1C).

**Fig. 1.**
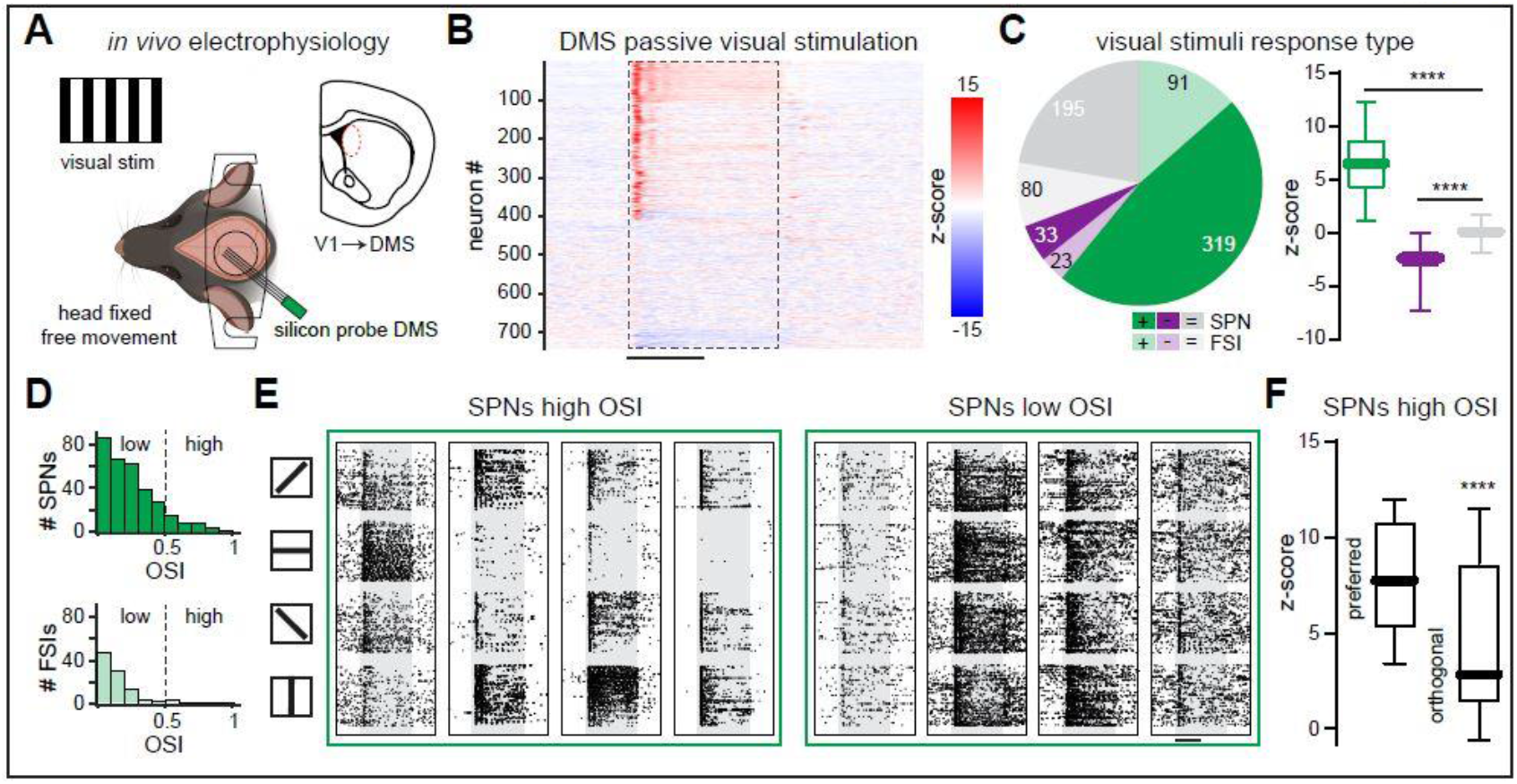
Responses of striatal neurons to visual stimuli. (**A**) Schematic representation of electrical recordings and visual stimuli in neurons from the dorsomedial striatum (DMS) receiving direct projections from primary visual cortex (V1) in awake head-fixed mice. (**B**) Z-scored activity profiles from 741 striatal neurons during passive visual stimuli with four different orientations recorded in 24 mice. Scale bar, 1 s. (**C**) Left: Quantification of striatal response types to visual stimuli of putative spiny projection neurons (SPNs) and fast spiking interneurons (FSIs). Right: increased responses to visual stimuli (green) and decreased responses to visual stimuli (purple) were significantly different from non-responsive neurons (gray). ^****^ P < 0.0001; Kruskal Wallis test Multiple comparisons; increased responses: 410 neurons; decreased responses: 56 neurons; non-responsive: 275 neurons. (**D**) Histogram of orientation selectivity index (OSI) for SPNs (top) and FSIs (bottom). (**E**) Representative responses to different orientations of drifting-gratings from SPNs with high OSI and SPNs with low OSI. Each column shows a different SPN. Scale bar, 1s. (**F**) Visal responses to the preferred orientation of SPNs with high OSI were higher compared to the orthogonal orientation. ^****^ P < 0.0001; Mann-Whitney test; n = 36 neurons. Data in (**C**) and (**F**) are presented as box-and-whisker plots displaying median, interquartile, and range values.

V1 projections to the DMS can evoke excitatory responses in striatal neurons (*22*) suggesting that DMS may process basic visual features. We found that, among SPNs responding to drifting-gratings, 36/319 showed high orientation selectivity index (OSI) while the rest exhibited low OSI. In contrast, only 2/91 FSIs that responded to visual stimulation displayed high OSI (Figure 1D), indicating that striatal FSIs are broadly tuned to orientation as has also been observed in V1 (*5*). SPNs with high OSI showed increased firing rates for their preferred orientation compared to a suppression in firing for their orthogonal orientation (Figure 1, E and F), whereas SPNs with low OSI showed increased responses to different orientations (Figure 1E). Our results show that a subset SPNs located in a target region of V1 processes basic visual information during passive visual stimulation, without relying on behavioral conditioning.

### Striatal population activity reliably predicts specific features of visual stimuli

We next examined whether striatal population activity could predict different features of visual stimuli. To do so we isolated population responses of SPNs to visual stimuli with different orientations and observed distinctive activity profiles (Figure 2A), suggesting the existence of specialized striatal ensembles for different orientations. Interestingly, we observed that SPNs with low OSI showed increased responses to their preferred orientation compared to their orthogonal orientation (Figure 2B) resembling SPNs with high OSI (Figure 1F). Applying demixed principal component analysis (dPCA) (*29*) to SPNs that increased their responses to visual stimuli, we identified clear orientation-specific population trajectories defined by all responsive SPNs (Figure 2C). Notably, orientation-specific population trajectories persisted even when only SPNs with low OSI were considered (Figure 2D). To measure the accuracy of striatal population activity to predict the orientation of drifting-gratings we used support vector classifier (SVC) models. We observed that even SPNs with low OSI reliably predicted specific features of visual stimuli (Figure 2E). The observation that SPNs with low OSI can predict the orientation of drifting-gratings similarly to the prediction of all responsive SPNs suggests that striatal feature selectivity not only emerges from high OSI SPNs but also from the functional organization of intra-striatal circuits.

**Fig. 2.**
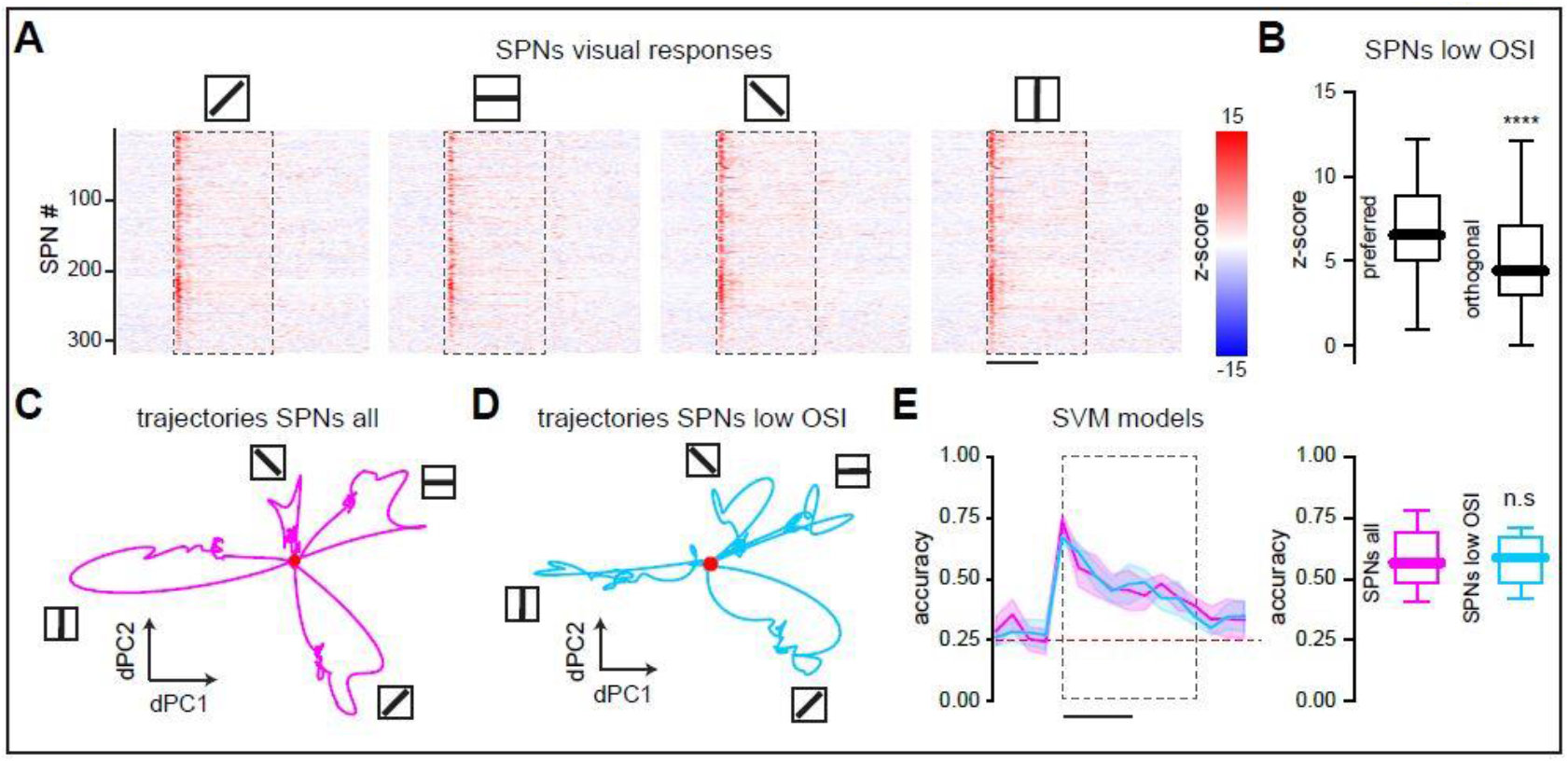
SPNs with low OSI predict different features of visual stimuli. (**A**) Z-scored activity profiles from 319 SPNs that increased their responses to different orientations of drifting-gratings. Scale bar, 1s. (**B**) Visal responses to the preferred orientation of SPNs with low OSI were higher compared to the orthogonal orientation. ^****^ P < 0.0001; Mann-Whitney test; n = 280 neurons. (**C**) Two-dimensional representation of neuronal population trajectories of all SPNs that increased their responses to visual stimuli with different orientations computed with demixed principal component analysis (dPCA). Note distinctive trajectories for different orientations of visual stimuli. Red dot indicates the starting point of the trajectory (visual stimuli onset). (**D**) Distinctive neuronal population trajectories for different orientations of visual stimuli are also observed for SPNs with low OSI. Red dot indicates the starting point of the trajectory. (**E**) The accuracy of support vector classifier (SVC) models trained with all responsive SPNs (high OSI and low OSI; magenta) is similar to the accuracy of SVM models trained only with low OSI SPNs (blue). n. s. P = 0.9626; Mann-Whitney test; n = 20 points. Data in (**B**) and (**E**) are presented as box-and-whisker plots displaying median, interquartile, and range values.

### Striatal spiny projection neurons from the direct and indirect pathways encode features of visual stimuli

To investigate the functional organization of striatal circuits involved in visual processing, we analyzed visually evoked responses across subpopulations of SPNs. To do so, D1-Cre or D2-Cre mice received striatal injections of AAV-EF1a-DIO-hChR2(H134R)-EYFP, enabling Cre-dependent expression of channelrhodopsin-2 in the DMS (Figure 3A). We achieved cell-type-specific identification of direct pathway (dSPNs) or indirect pathway (iSPNs) neurons by optotagging SPNs responding to blue light (Figure 3B). We observed that subpopulations of optotagged SPNs from both the direct and indirect pathways have similar responses to optogenetic stimuli (Figure 3C). Optotagged dSPNs and optotagged iSPNs reliably responded to visual stimuli (Figure 3D). Moreover, responses to visual stimuli (Figure 3E) and visual response latencies (Figure 3F) were similar between dSPNs and iSPNs. From optotagged SPNs that responded to visual stimuli, we identified dSPNs and iSPNs with high OSI as well as dSPNs and iSPNs with low OSI. Orientation selectivity between dSPNs and iSPNs was not significantly different (Figure 3G). These results suggest parallel contributions to visual feature encoding from both striatal pathways.

**Fig. 3.**
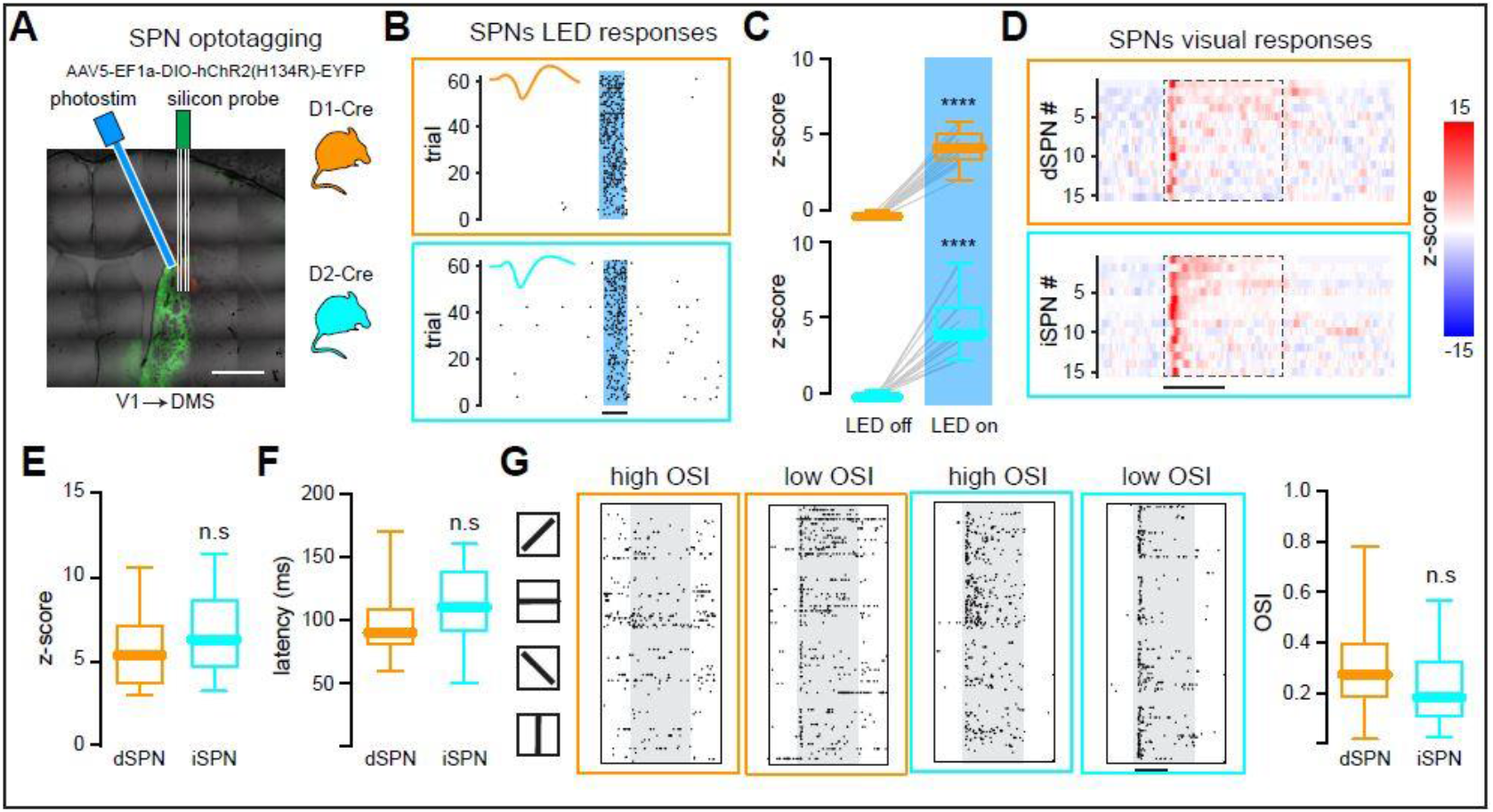
Striatal projection neurons from the direct and indirect pathways participate in visual feature encoding. (**A**) Schematic representation of electrical recordings and optotagging. Optogenetic photostimulation was delivered through a fiber optic attached to a blue LED positioned next to the recording probe. Image shows EYFP fluorescence of DMS expressing ChR2. Scale bar, 500 µm. D1-Cre mice (orange) or D2-Cre mice (cyan) were used to identify neurons from the direct pathway (dSPNs) or neurons from the indirect pathway (iSPNs), respectively. (**B**) Optogenetic evoked responses to blue light (100 ms) from a representative dSPN (top; orange) and a representative iSPN (bottom; cyan). (**C**) Optotagged neurons were identified by increased responses to blue light (LED on). Top: ^****^ P < 0.0001; Wilcoxon matched-pairs signed rank test; n = 15 dSPNs from 6 mice. Bottom: ^****^ P < 0.0001; Wilcoxon matched-pairs signed rank test; n = 15 iSPNs from 6 mice. (**D**) Z-scored activity profiles of optotagged dSPNs (top, orange) and optotagged iSPNs (bottom, cyan) that also responded to visual stimuli. Scale bar, 1 s. (**E**) dSPNs and iSPNs had similar responses to visual stimuli. n.s. P = 0.1736; Mann-Whitney test; n = 15 dSPNs from 6 mice; n = 15 iSPNs from 6 mice. (**F**) dSPNs and iSPNs had similar response latencies to visual stimuli. n.s. P = 0.1468; Mann-Whitney test; n = 15 dSPNs from 6 mice; n = 15 iSPNs from 6 mice. (**G**) Representative responses to different orientations of drifting-gratings from optotagged SPNs with high OSI and optotagged SPNs with low OSI. Each column shows a different SPN. Scale bar, 1s. OSI was not different between dSPNs and iSPNs. n.s. P = 0.3046; Mann-Whitney test; n = 15 dSPNs from 6 mice; n = 15 iSPNs from 6 mice. Data (**C**), (**D**), (**F**), and (**G**) are presented as box-and-whisker plots displaying median, interquartile, and range values. Lines in (**C**) show individual neurons.

### Striatal visual responses depend on intra-striatal connectivity between spiny projection neurons

To further characterize how intra-striatal connectivity between dSPNs and iSPNs influences visual processing, we used Cre-dependent Designer Receptors Exclusively Activated by Designer Drugs (DREADD) expression to selectively silence either dSPNs in D1-Cre mice or iSPNs in D2-Cre mice while recording striatal visual responses (Figure 4A). Selective silencing of either pathway inhibited striatal responses to visual stimuli in a subpopulation of SPNs but disinhibited visual responses in a different subpopulation of SPNs (Figure 4B) suggesting that intra-striatal connectivity participates in visual feature extraction. The overall activity of inhibited and disinhibited subpopulations of SPNs remained balanced before and after selective silencing (Figure 4C). Remarkably, selective silencing of either pathway reduced high orientation selectivity in SPNs (Figure 4D) indicating that intra-striatal connectivity between dSPNs and iSPNs is necessary to reliably encode features of visual stimuli. Our results demonstrate that striatal functional modules for visual information processing require the activation of dSPNs and iSPNs challenging the classical view of antagonistic activation patterns between dSPNs and iSPNs (*30, 31*). Moreover, considering known intra-striatal connectivity patterns (*32, 33*) we propose a novel striatal functional module comprised by dSPNs and iSPNs that encode the same visual features conveyed by specialized cortical ensembles (Figure 4E). Thus, intra-striatal connectivity and activation of both dSPNs and iSPNs take part in feature-specific extraction during visual information processing.

**Fig. 4.**
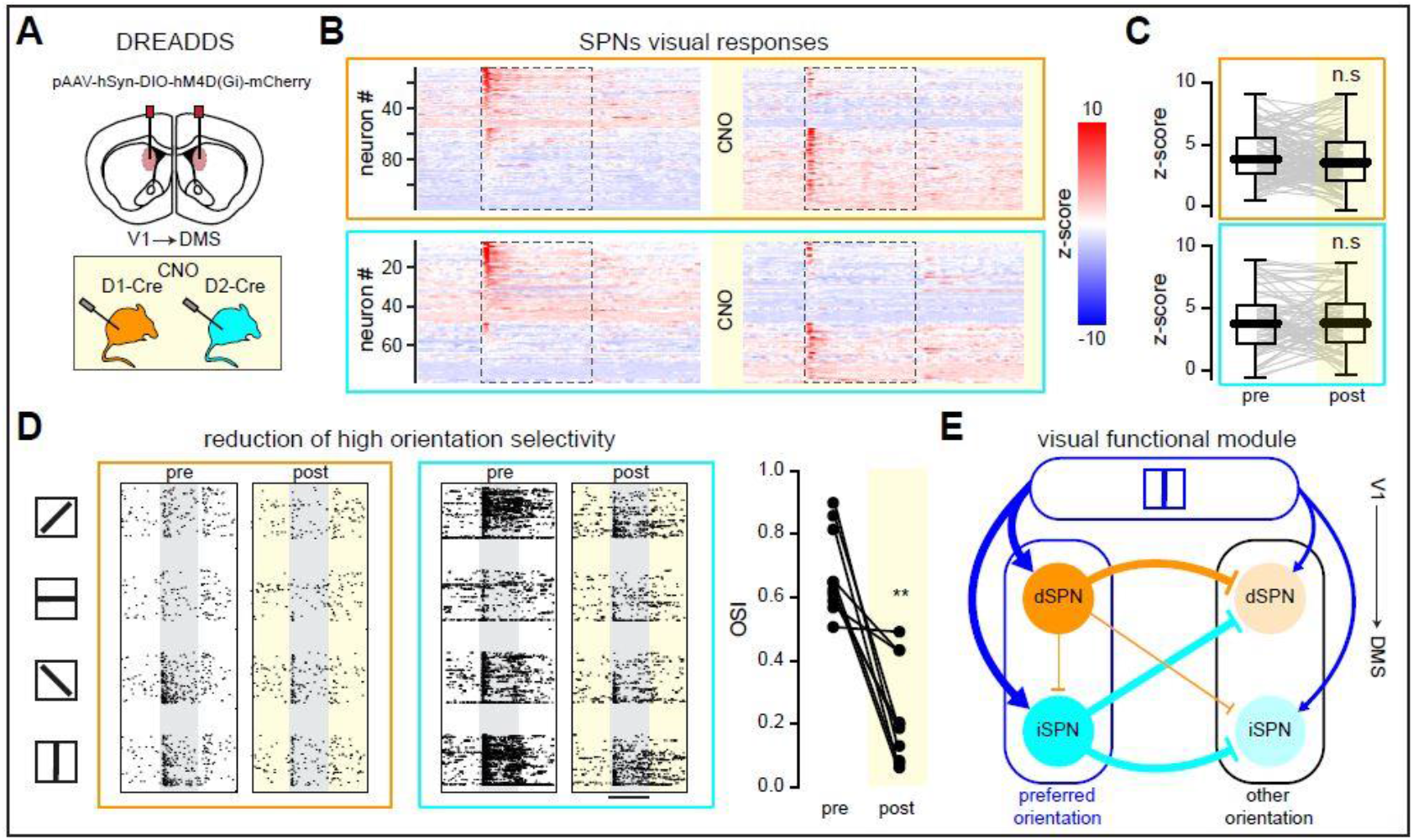
Striatal visual information processing depends on intra-striatal connectivity. (**A**) Schematic representation of Designer Receptors Exclusively Activated by Designer Drugs (DREADDS) injected bilaterally in the dorsomedial striatum (DMS) of D1-Cre (orange) or D2-Cre mice (cyan). (**B**) Z-scored activity profiles from SPNs that responded to visual stimuli either before or after CNO application in D1-Cre (top; orange) or D2-Cre (bottom; cyan) mice. Note that a subpopulation of SPNs is inhibited by the specific silencing of the direct or the indirect striatal pathways, whereas another subpopulation of SPNs is released. (**C**) Responses to visual stimuli remained balanced after the selective silencing of dSPNs (top; orange; n.s. P = 0.1120; Wilcoxon matched-pairs signed rank test; n = 111 SPNs from 6 mice) or iSPNs (bottom; cyan; n.s. P = 0.7889; Wilcoxon matched-pairs signed rank test; n = 73 SPNs from 5 mice). (**D**) Representative responses to different orientations of drifting-gratings before and after CNO from SPNs with high OSI in D1-Cre (orange) or D2-Cre (cyan) mice. High OSI was reduced after CNO (^**^ P = 0.0020; Wilcoxon matched-pairs signed rank test; n=10 SPNs from 11 mice). (**E**) Proposed striatal functional module for visual information processing highlighting dSPNs (orange) and iSPNs (cyan). Line widths represent the strength of cortical projections (blue) or intra-striatal axon collaterals between SPNs (orange and cyan). Cortical neurons encoding the preferred orientation strongly connect to both dSPNs and iSPNs, whereas the same cortical neurons weakly target dSPNs and iSPNs that encode other orientations.

## Discussion

Combining publicly available cortico-striatal connectivity mapping with chemogenetics and opto-targeted population recordings, we identified a striatal circuit that processes basic visual features during passive visual stimulation. Unlike traditional striatal studies that rely on conditioned behaviors linked to visual cues, our approach reveals encoding of visual stimuli by striatal neurons in the absence of behavioral contingencies. The involvement of striatal inhibitory circuits in visual processing challenges the prevailing cortico-centric hierarchy for visual perception, suggesting novel principles of mammalian visual system organization.

### Organization of striatal functional modules

In primary visual cortex, it has been demonstrated that different visual stimuli are encoded by distinctive cortical neurons (*3, 4*). Our findings reveal that striatal neurons also encode visual stimuli, establishing both a novel experimental framework and conceptual paradigm for understanding striatal functional organization in visual processing beyond cortical circuits. The description of striatal functional modules has been elusive up to date due to difficulties distinguishing between motivation, motor, learning, and sensory signals. It has been proposed that intra-striatal lateral inhibition shapes striatal functional modules where dSPNs and iSPNs have antagonistic activity patterns (*30, 31*). However, because experimental evidence comes from movement related striatal areas it has been difficult to dissect the basic functional organization of striatal modules. In our experiments, striatal visual responses were analyzed while mice remained still, enabling us to characterize a striatal circuit for visual processing detached from locomotion-related activity. We observed activation of both dSPNs and iSPNs for the same visual orientation indicating that both striatal pathways are part of the same striatal functional module. Furthermore, our results revealed that striatal orientation selectivity requires intra-striatal connectivity between the direct and indirect BG pathways. These findings are consistent with established models of striatal population activity mediated by local inhibition (*30, 34*), and suggest that the DMS perform additional processing of specific visual information features conveyed by cortical projections, redefining the current paradigm of striatal functional modules.

### A novel cortico-striatal circuit for visual information processing

It has been suggested that striatal activity is mainly related to action selection, locomotion, movement kinematics, and behavioral conditioning (*9, 31, 35, 36*). However, we observed that features of passive visual stimuli are encoded by striatal neurons, indicating that cortico-subcortical circuits participate in visual perception. Our results agree with recently published data showing that under passive visual stimulation, reliable neuronal responses can be observed in primary visual cortex and DMS (*27*), suggesting a specialized cortico-striatal circuit for visual processing even in the absence of behavioral conditioning.

The balanced interaction between dSPNs and iSPNs during visual stimuli points out to the existence of specialized visual functional modules comprised by cortico-striatal subpopulations, expanding the classical view of BG models (*37–39*) that excludes a basal-ganglia-thalamocortical circuit that processes basic features of visual stimuli. It has been shown in mice that activation of the direct striatal pathway enhances cortical activity whereas activation of the indirect striatal pathway decreases cortical activity (*40*) indicating that striatal functional modules comprised by dSPNs and iSPNs could refine cortical activity related to specific features of visual stimuli. Moreover, a basal-ganglia-thalamocortical loop originating from V1 that comprises both the direct and indirect striatal pathways has been anatomically described (*41*). Accordingly, our results suggest the existence of a specialized basal ganglia-thalamocortical circuit that processes visual information in parallel to other brain functions.

### Relevance of visual information processing by striatal neurons

It has been proposed that different portions of the dorsal striatum are engaged during the early and late phases of motor skill learning. Thus, the DMS has been involved in the initial stages of motor learning representing goal-directed actions, whereas the DLS has been related to habit formation (*7, 11*). Different from most studies of DMS function that have focused on learning to associate visual cues with stereotyped movements (*9*), our results propose a visuospatial function of the DMS. It has been shown that this part of the DMS receives direct projections from primary, posteromedial, and lateral visual cortex. Moreover, it also receives direct projections from the medial entorhinal cortex and the dorsal, ventral, and agranular retrosplenial areas (*21*) suggesting that striatal functional modules may process specific features of the environment. Interestingly, layer 2/3 and layer 5 neurons of V1 project directly to DMS (*22*) indicating that specific aspects of visual stimuli are distributed in parallel to higher cortical visual areas and subcortical structures such as if a given feature of visual stimuli can be processed by parallel distributed circuits for different functions. Thus, striatal functional modules described here represent a visual window to the external world that could be involved in visual perception. This is supported by recent data demonstrating that the overall activity in each part of the striatum closely mirrors the activity of anatomically associated cortical regions regardless of task engagement (*42*) highlighting that visual cortical features are preserved in their corresponding striatal structures.

Our results provide a starting point to understand the role of basal ganglia structures in visual perception, opening the door for the development of novel treatments for neurological conditions where abnormal visual processing is common such as Parkinson’s disease (*43, 44*), Schizophrenia (*45*), or Autism spectrum disorder (*46*). Based on our findings we propose that current models of visual perception should incorporate contributions from basal ganglia-thalamocortical circuits.

## Acknowledgements

We thank Deisy Gasca Martinez, Ericka A. de los Rios Arellano, Martín Garcia Servin, Alejandra Castilla Leon, Eugenia Ramos Aguilar, Adriana Gonzalez Gallardo, Anaid Antaramian Salas, Maria A. Carbajo Mata, and Elsa Nydia Hernández Rios for technical assistance.

## Author Contributions

L. C-R. and J. P-B. designed research; J. P-B. and LA. T performed experiments; J. P-B. and R. V-C. analyzed data; L. C-R. wrote the manuscript.

## Data availability

All data is stored in the Functional Circuits Reprogramming Laboratory and will be made available upon reasonable request.

## Materials and Methods

### Animals

All experimental procedures were carried out in accordance with the guidelines of the Bioethics Committee of the Neurobiology Institute for the care and use of laboratory animals that comply with the standards outlined by the Guide for the Care and Use of Laboratory Animals (NIH) and the Policies on the Use of Animals in Neuroscience Research. Experiments were performed on C57BL/6, D1-dopamine receptor Cre-recombinase [D1-Cre, Tg(Drd1-cre)EY262Gsat, Gensat], or D2-dopamine receptor Cre-recombinase [D2-Cre, B6.FVB(Cg)-Tg(Drd2-cre)ER44Gsat/Mmucd, Gensat] male mice at postnatal day 60-120. Animals were housed on a 12h light-dark cycle with food and water ad libitum.

### Headplate implantation and viral injections

Mice were anesthetized with isoflurane (1-2%) and placed in a stereotaxic system (Stoelting). All procedures were performed in sterile conditions. During the surgery, eyes were moisturized with eye ointment. Respiratory rate and tail pinch reflex were monitored along the surgery. A custom designed stainless steel head plate was attached to the skull using dental cement (C&B Metabond). Additionally, two metal pins were implanted on the surface of the brain to serve as reference and ground. After surgery, mice received carprofen injections (5 mg/kg) for 2 days as post-operative pain medication. Then, mice were handed and exposed to head fixation conditions for 4 days before experiments. Experiments were performed 1 week after headplate implantation.

To identify striatal neurons belonging to the direct or indirect pathways, 600 nL of the Cre-dependent viral construct AAV5-EF1a-DIO-hChR2(H134R)-EYFP (Addgene) were unilaterally injected into the DMS (AP: 0.5 mm; ML: −1.4 mm; DV: −3 mm) of D1-Cre or D2-Cre anesthetized mice (isoflurane 1-2%) at a rate of 50 nL/min using a Hamilton 1.0 µL Neuros Model 7001KH syringe mounted on a Pump 11 Elite (Harvard Apparatus). Channelrhodopsin-2 viral injections were performed in a stereotaxic system (Stoelting) in the DMS contralateral to the eye where visual stimuli were presented. To ensure proper viral diffusion, the needle was left in place for 10 minutes before being slowly withdrawn. Viral injection was performed 4 weeks before headplate implantation.

To selectively silence SPNs from the direct or indirect pathways, 400 nL of the Cre-dependent viral construct pAAV-hSyn-DIO-hM4D(Gi)-mCherry (Addgene) were injected bilaterally into the DMS of D1-Cre or D2-Cre anesthetized mice (isoflurane 1-2%) as described above. Viral injection was performed 4 weeks before headplate implantation.

### Electrophysiology

For electrophysiological recordings a 2mm diameter craniotomy was made above the DMS of anesthetized mice (isoflurane 1-2%) 6 days after headplate implantation. The exposed brain was protected with a biocompatible silicone cap (Kwik-Sil) before the experiments. One day after the craniotomy, mice were head-fixed on a custom-designed treadmill. Electrophysiological signals were acquired with silicon probes (NeuroNexus, 64-channels, A4 x A4 tet-5mm), inserted vertically in the DMS (AP: 0.5 mm; ML: −1.4 mm; DV: −2.5 mm), contralateral to the eye where visual stimuli were presented, using OmniPlex Neural recording data acquisition system (Plexon) at a sampling rate of 40 kHz. The exposed brain was regularly irrigated with Dulbecco’s phosphate-buffered saline (DPBS, Sigma Aldrich) to prevent tissue dehydration. During recording sessions mice were awake and allowed to move freely on the treadmill.

### Visual stimulation

Visual stimuli were generated using Psychophysics Toolbox (MATLAB) and displayed on a monitor distanced 15 cm from the right eye at 45° to the middle axis of the head. Different drifting-gratings were presented for 2 s, with inter-stimulus interval of 8 seconds. During recording sessions each direction was randomly presented for 45 trials. Full-field drifting-gratings (contrast: 100%, 0.035 cycles/°, 2 cycles/sec) consisted in four different orientations (0°, 45°, 90°, 135°). In the absence of visual stimuli, the monitor displayed a gray screen with mean luminescence similar to visual stimuli. Locomotion was measured with an angular position magnetic sensor attached to the treadmill. Electrophysiological recordings were synchronized with visual stimuli and locomotion using a Digital Acquisition Board (Arduino Uno) connected to a host computer.

### Classification of visually evoked activity

To evaluate the accuracy of striatal population activity to classify different orientations of drifting-gratings support vector classifier (SVC) models were trained using SPNs that responded to visual stimuli. The regularization parameter was se to 0.8. To measure the accuracy of SVC models before and after visual stimuli recordings were partitioned into contiguous non-overlapping time bins of 250 ms. The model architecture was preserved for different time points and different subsets of SPNs. A 5-fold-cross-validation procedure was used. Trials were divided into 5 blocks preserving the proportion of samples across all classification classes. For each iteration, four blocks were used for training and one for validation. Trials were randomly shuffled in each partition. The accuracy of the classifier was then computed for all SPNs that increased their responses to visual stimulation and for SPNs with low OSI. Then, four consecutive time windows (250 ms each) per fold were used to compared the accuracy of all SPNs against low OSI SPNs (20 accuracy values for each subpopulation of SPNs).

### Optotagging of striatal neurons from the direct or indirect pathways

For optotagging experiments D1-Cre or D2-Cre mice injected with channelrhodopsin-2 were head-fixed on a custom-designed treadmill one day after craniotomy for electrophysiological recordings. A fiber optic cannula of 400 µm diameter attached to a fiber optic and connected to a blue LED (470 nm) was inserted into the DMS through the craniotomy at a 20° angle from the vertical line. The tip of the fiber optic cannula was placed 300 µm away from the silicon probe. 60 light pulses (100 ms, 4 mW,) were irradiated using a LED controller (CD2100, Thorlabs). Neurons whose firing rate increased 3 times above their baseline activity within the first 40 ms of blue light pulses in D1-Cre or D2-Cre channelrhodopsin-2 expressing mice were considered as belonging to the direct (dSPNs) or indirect (iSPNs) pathways respectively.

### Selective silencing of striatal neurons from the direct or indirect pathways

For chemogenetic experiments D1-Cre or D2-Cre mice injected with inhibitory DREADDs were implanted, at the same time of headplate implantation procedures, with an intra-peritoneal catheter through a small midline incision into the abdomen where a purse-string suture was tightened around tubing subcutaneously tunneled, exteriorized, and secured on the back of the mice. This procedure was used to avoid stress and animal movement caused by intra-peritoneal injections. One day after craniotomy, mice were head-fixed on a custom-designed treadmill for electrophysiological recordings. Visual responses of the same neurons were recorded before and after CNO administration. CNO (0.5 mg/kg) was delivered intra-peritoneally through the tubing after the first protocol of visual stimuli presentation without moving the mice. 30 minutes after CNO administration, a second protocol of visual stimuli was presented to mice.

### Spike sorting

Extracellular electrophysiological recordings were processed with a high-pass filter above 350 Hz. Spiking events were detected on-line as signals crossing a threshold determined for each neuron. 5 ms waveforms samples were recorded before and after the threshold crossing. To isolate units from electrophysiological recordings, spike sorting was performed with a semi-automatic clustering procedure based on principal component analysis (PCA) and k-means using the software Plexon Offline Sorter. Spike sorting was performed on concatenated waveform traces from the four channels of each tetrode.

### Neuronal classification based on electrophysiological signals

Putative spiny projection neurons (SPNs) and putative fast spiking interneurons (FSIs) were classified according to the waveform with the highest amplitude of each tetrode array. Waveforms from each neuron were rescaled and aligned to calculate mean value. The mean waveform of each neuron was projected into a two-dimensional space using uniform manifold approximation and projection algorithm (UMAP). Using this approach, units were reliably separated into two main clusters depicting FSIs with narrow spiking or SPNs with broad spiking.

### Analysis of individual neuronal responses to visual stimuli

Responses for each visual stimuli were calculated binning the duration of visual stimuli within 100 ms sliding time windows with 10 ms overlap. Then, the baseline firing rate, calculated by averaging the firing rate over 1 second before visual stimuli presentation, was subtracted from each time window. To discard the effect of locomotion, only trials where mice remained still without running were considered for analysis.

According to their responses during the first 250 ms after visual stimuli onset, neurons were classified into three groups: 1. Neurons that increased their responses above three standard deviations from the baseline. 2. Neurons that were not responsive. 3. Neurons that decreased their responses below three standard deviations from the baseline.

The preferred orientation was determined by the drifting-grating that evoked the highest average firing rate for each neuron. Then, orientation selectivity index (OSI) was calculated as (Rpref – R_ortho_)/(R_pref_ + R_ortho_), where R_pref_ denotes the preferred orientation and R_ortho_ denotes the orthogonal orientation to R_pref_. Neurons with OSI>0.5 were considered highly selective.

### Population trajectory analysis

Demixed principal component analysis (dPCA) was used to factorize population activity into neuronal trajectories that reflect their dependence on different visual stimuli (*29*). A matrix X containing the average neuronal activity for each corresponding visual stimulus of all recorded mice for a given experiment was used. For each visual stimulus S there are a collection of K trials with N neurons in each trial and T time points. Neuronal population trajectories were constructed using the first two components with most variability explained by different visual stimuli. Different subpopulations of neurons were used for each of the trajectories shown in the figures. The resulting neural trajectories were smoothed using a Gaussian kernel of 200 ms. Distinctive population trajectories indicate that neuronal responses encode different features of visual stimuli.

### Histological procedures

At the end of experimental protocols, mice were deeply anesthetized (isoflurane 5%) and intracardially perfused with 4% paraformaldehyde in PBS. The brain was extracted and sectioned coronally in 100 µm slices using a cryostat. To visualize the recording sites, silicon probes were coated with DiI (Invitrogen). To visualize channelrhodopsin-2 or DREADDs viral infections, EYFP or mCherry were imaged on an epifluorescence microscope.

### Statistical analysis

Statistical power analysis to determine the number of animals used in each experiment was not used. Sample size was determined based on previous publications describing responses to visual stimuli in mice (*3–5*). Statistical test were done in GraphPad Prism. Statistical details for each experimental group can be found in figure legends. Data presented as whisker boxplots display median, interquartile, and range values. For electrophysiological experiments n refers to the number of neurons.

